# Size aftereffect is non-local

**DOI:** 10.1101/2020.03.19.998161

**Authors:** Ecem Altan, Huseyin Boyaci

## Abstract

It is well known that prolonged exposure to a certain size stimulus alters the perceived size of a subsequently presented stimulus at the same location. How the rest of the visual space is affected by this size adaptation, however, has not been systematically studied before. Here, to fill this gap in literature, we tested size adaptation at the adapter location as well as the rest of the visual space. We used peripherally presented solid discs (Experiment 1) and rings (Experiment 2) as adapter and target (test) stimuli. Observers adapted to a mid-sized stimulus and judged the size of the subsequently presented smaller or larger target stimuli. Results showed that the perceived sizes of target stimuli were repelled away from the adapter size, not only at the adapter location but also at distant locations. These findings demonstrate that size adaptation causes widespread distortion of the visual space and alters perceived size. We discuss possible computational models that may underpin the perceptual effect.

## 1 Introduction

Perceived size of an object strongly depends on contextual factors such as relative size of the surrounding stimuli (Ebbinghaus, 1902; Massaro & Anderson, 1971) and perceived depth (Zanforlin, 1967; Holway & Boring, 1941). Many well-known size illusions clearly demonstrate these effects (Murray, Boyaci, & Kersten, 2006; Fang, Boyaci, Kersten, & Murray, 2008). Furthermore, Blakemore and Sutton (1969) demonstrated that perceived size can also be affected by temporal context through adaptation. They found that after prolonged exposure to a high-contrast grating pattern of a certain spatial frequency, perceived frequency of a subsequently presented grating pattern shifts away from the adapted spatial frequency. More specifically, after a certain period of adaptation to a mid-level frequency grating, a higher (lower) frequency grating is perceived to have a higher (lower) frequency than its actual frequency. This *repulsive* perceptual shift (i.e. aftereffect) is a typical consequence of visual adaptation that can be observed with many different stimulus features such as orientation (Jin, Dragoi, Sur, & Seung, 2005), shape (Suzuki & Cavanagh, 1998), motion (Mather, Pavan, Campana, & Casco, 2008), and faces and facial expressions (Watson & Clifford, 2003; Yang, Hong, & Blake, 2010).

Compared to the number of studies on the effect of spatial context on size perception, there are relatively few studies on temporally-induced size adaptation. Recent such studies (e.g. Pooresmaeili, Arrighi, Biagi, & Morrone, 2013; Kreutzer, Fink, & Weidner, 2015; Laycock, Sherman, Sperandio, & Chouinard, 2017; Zeng, Kreutzer, Fink, & Weidner, 2017) have mainly addressed where in the visual pathway the size aftereffect emerges. But relatively few studies investigated the spatial aspects of the size adaptation (Kreutzer, Ralph, & Fink, 2015). Most critically, previous studies on size adaptation tested the perceptual effect always at the same visuospatial location as the adapter. Whether the perceived size can be distorted by a distant adapter remains unanswered. The answer to this question has the potential to provide critical information about size adaptation and its characteristics over space, which in turn could lead to a better understanding of the mechanisms involved in size perception. Moreover, it can potentially motivate other questions in size adaptation research, as well as in adaptation studies in general.

The current study addresses the aforementioned gap in the literature and aims to reveal the spatial extent of the size aftereffect. For that purpose, two behavioral experiments were conducted with two different types of stimuli. In both of these experiments, the effect of adaptation to a certain-sized circular stimulus was tested at multiple locations including the adapter’s location and other locations that do not have a recent stimulation history.

## 2 Experiment 1

### 2.1 Materials and Methods

#### 2.1.1 Participants

Three groups of twelve subjects participated in different parts of this experiment, at different times. First group (6 males, 6 females; age range: 22-32; *M* = 25.9; *SD* = 2.84) was tested for the center row locations in Figure 1.A, second group (5 males, 7 females; age range: 18-21; *M* = 19; *SD* = 1.13) was tested for the upper row locations, and third group (6 males, 6 females; age range: 18-26; *M* = 19.7; *SD* = 2.64) was tested for the lower row locations in the same figure. All subjects had normal or corrected-to-normal vision and gave their written informed consent prior to the experiment. Protocols and procedures were approved by the Bilkent University Human Ethics Committee.

**Figure 1:**
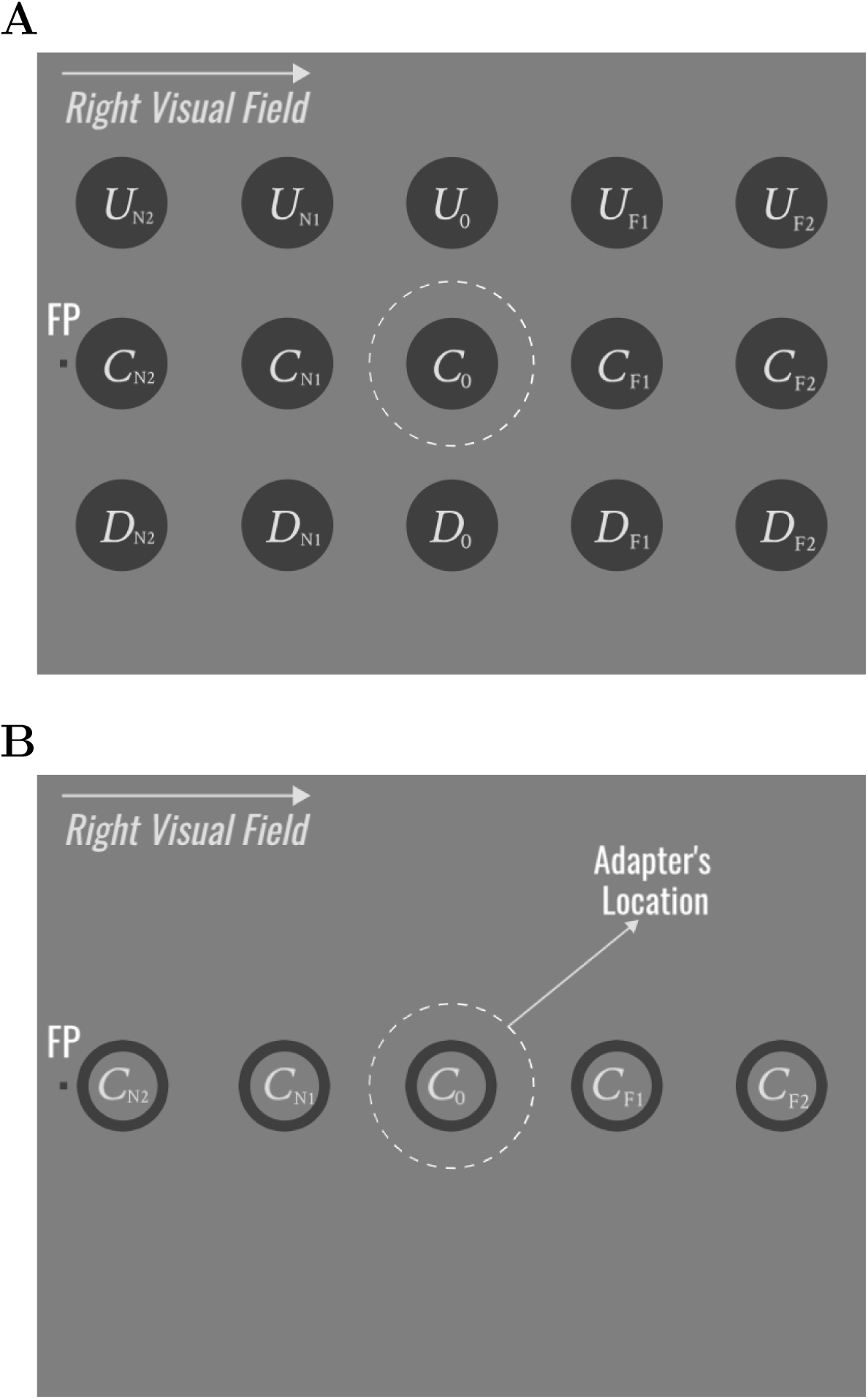
Spatial layout for Experiment 1 (A) and Experiment 2 (B). Dark gray discs (A) and rings (B) represent target stimuli and white dashed circles represent the size and position of the adapter (the adapter was a mid-sized disk in Experiment 1, a ring in Experiment 2). The adapter was presented 10.5° to the left or right of the fixation point (FP). Only the right visual field is shown here for clarity. The target stimuli appeared in only one of the shown positions in the test phase of a trial. The target could be either larger or smaller than the adapter. FP: fixation point, U: up, C: center, D: down, N2: nearer 2, N1: nearer 1, 0: zero, F1: further 1, F2: further 2.

#### 2.1.2 Stimuli and Apparatus

Stimuli were generated and presented using Psychophysics toolbox (Brainard, 1997) running on MATLAB (Mathworks). Participants were seated 65 cm away from a 30-inch NEC MultiSync LCD monitor (LCD3090WQXi; 60 Hz refresh rate; 1920 ×1200 screen resolution) in a dark room. A chin-rest was used to keep participants’ head stable.

A single experimental trial included an adaptation phase and a following test phase (Figure 2). Two phases were separated by a 300 ms interval. A mid-gray background was always present on the screen. Event duration and time intervals were determined based on studies in literature (e.g. Pooresmaeili et al., 2013).

**Figure 2:**
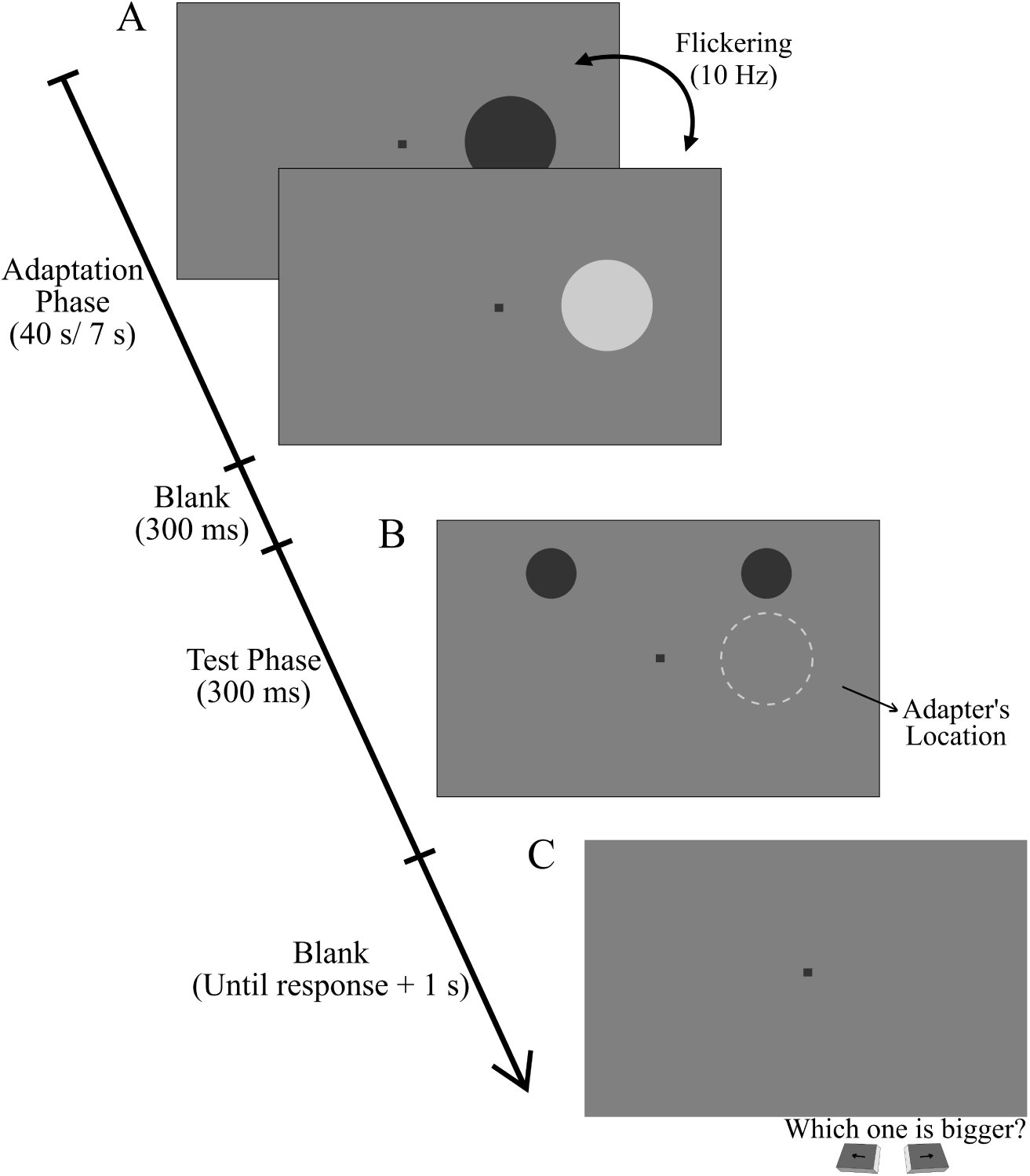
Time course of a single trial in Experiment 1. A. Trials started with an adaptation phase, in which a mid-sized flickering adapter was presented either in the left or the right visual field (in this case in right visual field; adapter hemifield was blocked in different sessions). Adaptation phase lasted 40s in the first trial and 7s in the remaining trials. This was followed by a 300ms blank fixation screen (same as C). B. Test phase lasted 300ms with a target disc (larger or smaller than the adapter; blocked in different sessions) in one of the fifteen positions in the adapted visual field, and a variable disc at the non-adapted visual field (the dotted line outlining the adapter location was not presented to the participants). The size of the variable disc in a trial was controlled by the experimental program based on the participant’s previous responses following an adaptive procedure. C. After the test phase, participants were required to press a key to indicate the bigger of the two discs (left or right). A final blank fixation screen remained for 1s after the participants responded. In the control blocks the adaptation phase was omitted.

##### Adaptation

In the adaptation phase, a mid-sized (diameter 2.5°) adapter disc was presented for 40s in the first trial in a block (initial adaptation), and for 7s in the remaining trials (top-up adaptation). In an experimental block, the adapter was presented only in the left or only in the right visual field, located at 10.5° away from the fixation point on the horizontal meridian. The adapter flickered at 10 Hz (from dark gray to light gray) in order to prevent afterimages.

##### Test

Two dark gray discs were presented in the test phase: A target disc was presented always at the same visual hemifield as the adapter, and a variable disc was presented always at the non-adapted visual hemifield. Target disc had always a fixed size of either 1.5° or 3.5° (smaller or larger compared to the adapter). The target sizes were determined based on our pilots with the intention of obtaining the maximum possible adaptation effect. As shown in Figure 1.A, target discs appeared at fifteen different locations in total. Three rows of test positions were named as up (U), down (D) and center (C); and five columns were named as nearer 2 (N2), nearer 1 (N1), zero (0), further 1 (F1) and further 2 (F2). The distance between each nearest neighbour position was 4°. The position of variable disc was always symmetrical to that of target disc with respect to the vertical center of the screen. The size of the variable was subject to an adaptive staircase procedure. The test phase lasted for 300ms.

#### 2.1.3 Procedure and Data Analysis

Participants started the experiment with a key press after reading the instructions on the introduction screen. They were required to fixate the mark at the center of the screen throughout the experiment (FP, fixation point). Following the adaptation and test phases, participants pressed one of the two arrow keys to indicate which disk appeared bigger in the test phase (right or left), while maintaining their fixation at the center on the blank screen. All trials followed the same event sequence as shown in Figure 2. The response of the participant was used to update the size of the variable disc for the following trials, using a 1-up 1-down staircase procedure with two interleaved staircases. Step size of the staircases was set to 0.32° in diameter at the beginning, but decreased by half after each response reversal, until it reached to 0.04°. Staircases had 25 trials each, which was carefully determined based on our pilots. Two staircases, one starting from a relatively bigger, the other from relatively smaller variable disc sizes, were used. This provided 50 trials for a single target location. A single block of experiment consisted of a total of 250 trials for all five target locations. Targets in five locations and from the two staircases were presented in a random order. Note that each participant group completed the tests in a single row shown in Figure 1.A.

Small and large targets, and left and right visual field adaptation were tested in separate adaptation blocks with a random order for each participant (4 conditions). In addition, participants completed 4 control blocks for the same 4 conditions without adaptation. Each session started with a control block and continued with an adaptation block after a short break. The target size and the visual hemifield in the adaptation and control blocks were determined randomly and independently. Participants completed the experiment in 4 sessions. The time interval between the sessions ranged from 2 hours to several days.

Point of subjective equality (PSE) for each target point and participant was computed by fitting a logistic function to the participant’s responses and determining the 50% point using the Psignifit 4 toolbox (Schütt, Harmeling, Macke, & Wichmann, 2016) on MATLAB. A PSE value corresponds to the size of the variable disc that is perceptually equal to that of the target disc for the participant.

Next, to quantify the size adaptation and apply further statistical tests, we computed an adaptation index.

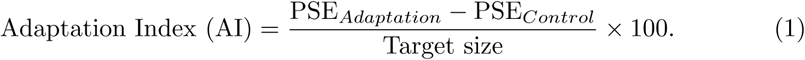

Negative AI values indicate perceptual underestimation, and positive values indicate perceptual overestimation of the target size after adaptation (*i*.*e*. smaller and larger perceived size compared to control respectively). Statistical analyses were performed on the AI values, using the JASP software (JASP Team, 2018). Since we had a strong a priori knowledge about the direction of the size aftereffects, we performed single tailed one sample *t* tests (*AI* < 0 for small target size; *AI* > 0 for large target size). Then a mixed ANOVA was performed to find whether there is any effect of visual field, and target positions (both within and between-subject factors) on AI.

### 2.2 Results

Figure 3 shows the adaptation index (AI) across the adapted visual hemifield. These maps clearly show that the adaptation effect is not limited to the location of the adapter but reaches distant positions in the visual field. Visual inspection of the maps also suggest that the adaptation effect on smaller targets is usually stronger than that on larger targets. Furthermore, the effect gets weaker as the distance to the adapter increases.

**Figure 3:**
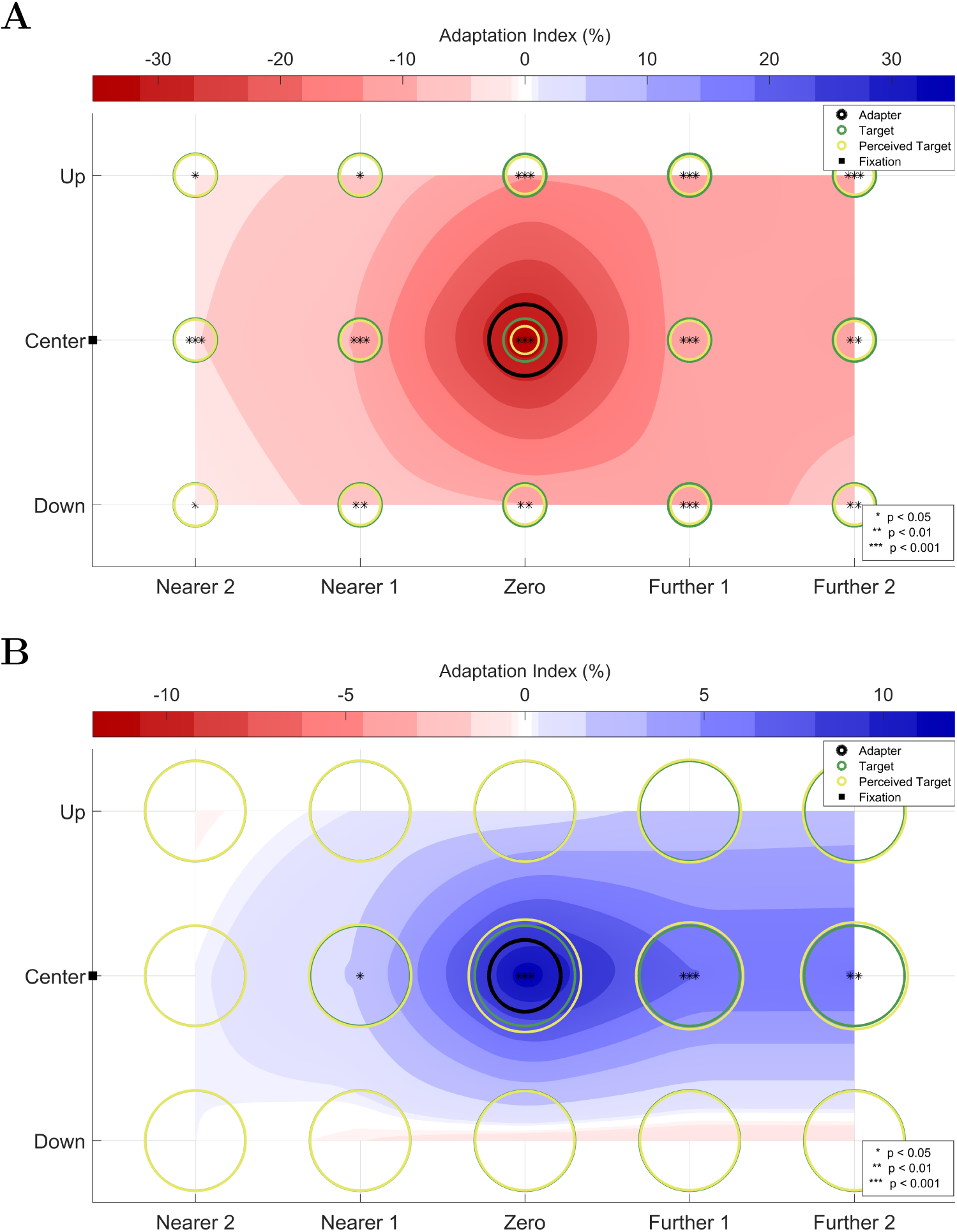
Adaptation effect for small (A) and large (B) target discs. To quantify the size adaptation effect, we computed an adaptation index (AI, see Equation 1). The maps were generated by averaging the AIs in left and right visual fields, as the effect was in the same direction regardless of the visual hemifield. Negative AI values (perceptual underestimation) are represented by shades of red, whereas positive AI values (perceptual overestimation) are represented by shades of blue. Darker shades mean greater adaptation effect. Note that the colormaps have different ranges in (A) and (B). AI values between the 15 target locations were estimated using natural neighbour interpolation. Thick black circles at the center-zero position show the relative size and position of the adapter. Thin green circles represent the actual sizes of targets and the yellow circles show their perceived sizes (all drawn to scale). Small black squares, next to ‘Center’ on the vertical axes, represent the fixation point. *p* values are FDR corrected.

On average (averaged across left and right visual fields) the small target disc presented at C_0_ position (i.e. concentric with the adapter) was perceived 35% (SEM=1.5%) smaller in the adaptation blocks, compared to the control blocks. Relatively weaker underestimations were observed in the remaining 14 eccentric positions. One sample *t* tests revealed that the AI values for small targets were significantly smaller than zero, at all of the fifteen positions (FDR corrected *p*s < 0.05; see Figure 3.A). Moreover, within subjects effect of a mixed ANOVA revealed a significant main effect of the adapter hemifield (*F*(1, 33) = 9.27, *p* < 0.01). Pairwise comparison for the adapter hemifield revealed stronger adaptation effect in the left visual hemifield than that in the right for small targets (*p* < 0.01). There was also a significant main effect of horizontal target positions (Sphericity was corrected via Greenhouse-Geisser correction; (*F*(2.62, 86.52) = 36.32, *p* < 0.001)) and significant between-subjects effect of vertical target positions (*F*(2, 33) = 7.86, *p* < 0.01). The interaction between the horizontal and the vertical positions was also significant (Sphericity corrected; *F*(5.24, 86.52) = 13.08, *p* < 0.001). Therefore, we performed simple main effect analysis and found that the difference among the levels of vertical positions was only significant at position Zero of the horizontal target positions (*p* < 0.001).

Large targets presented at C_0_ were perceptually 12% (SEM=1.7%) larger in the adaptation condition, compared to the control condition (averaged across two visual hemifields). In 9 out of 14 distant positions, the large target appeared larger in the adaptation condition compared to control condition. Analyses showed that the AI values were significantly larger than zero in 4 out of 15 positions (FDR corrected *p*s < 0.05; see Figure 3.B). A mixed ANOVA showed that there was a significant main effect of adapter region (*F*(1, 33) = 12.14, *p* < 0.01). The adaptation effect was significantly stronger in the left visual field, compared to that in the right visual field (*p* < 0.01). There was also a significant main effect of horizontal target positions (Sphericity corrected; *F*(2.69, 88.79) = 13.16, *p* < 0.001), a significant effect of vertical target positions (*F*(2, 33) = 14.23, *p* < 0.001); and a significant interaction between these two factors (Sphericity corrected; *F*(5.38, 88.79) = 13.64, *p* < 0.001). Simple main effect analysis showed that the difference among the levels of vertical positions was significant at the following positions: zero (*p* < 0.001), further 1 (*p* < 0.001) and further 2 (*p* < 0.01).

## 3 Experiment 2

In the second experiment we used rings instead of solid discs. We had two reasons for this choice. Firstly, we wanted to test whether the results of the first experiment would generalize to different types of stimuli. Secondly, we wanted the results of Experiment 2 to inform us about possible computational models explaining the size adaptation phenomenon. Because, whereas the solid adapter discs presumably activated and thus caused adaptation in a population of neurons whose receptive fields corresponded to the whole of the solid discs, the rings stimulated only the neurons receiving input from the thin lines that would correspond approximately to the boundaries of the disks. Thus, we envisaged that comparing adaptation effects across the two experiments could help us conceptualize computational models.

## 3.1 Materials and Methods

### 3.1.1 Participants

12 volunteers (4 males, 8 females; age range: 22-33; *M* = 25.1; *SD* = 2.97) participated in the experiment after giving their written informed consent. All participants had normal or corrected-to-normal vision. Protocols and procedures were approved by Bilkent University Human Ethics Committee.

### 3.1.2 Stimuli and Apparatus

Physical conditions were exactly the same as those in the previous experiment. Experimental design was also the same with few exceptions. First, both the flickering adapter and the target stimuli were rings with a linewidth of 0.18°, instead of discs. Second, only 5 target positions along the horizontal meridian were tested as opposed to the 15 positions in the first experiment (See Figure 1.B). Sizes and positions of rings matched exactly those of discs in the previous experiment.

### 3.1.3 Procedure and Data Analysis

Procedure and the data analyses were the same as in Experiment 1.

## 3.2 Results

Figure 4 shows the adaptation indices (AIs) for all tested target positions in Experiment 2, along with those from corresponding positions in Experiment 1. Clearly the adaptation effect is present and the pattern is nearly identical to that found in Experiment 1.

**Figure 4:**
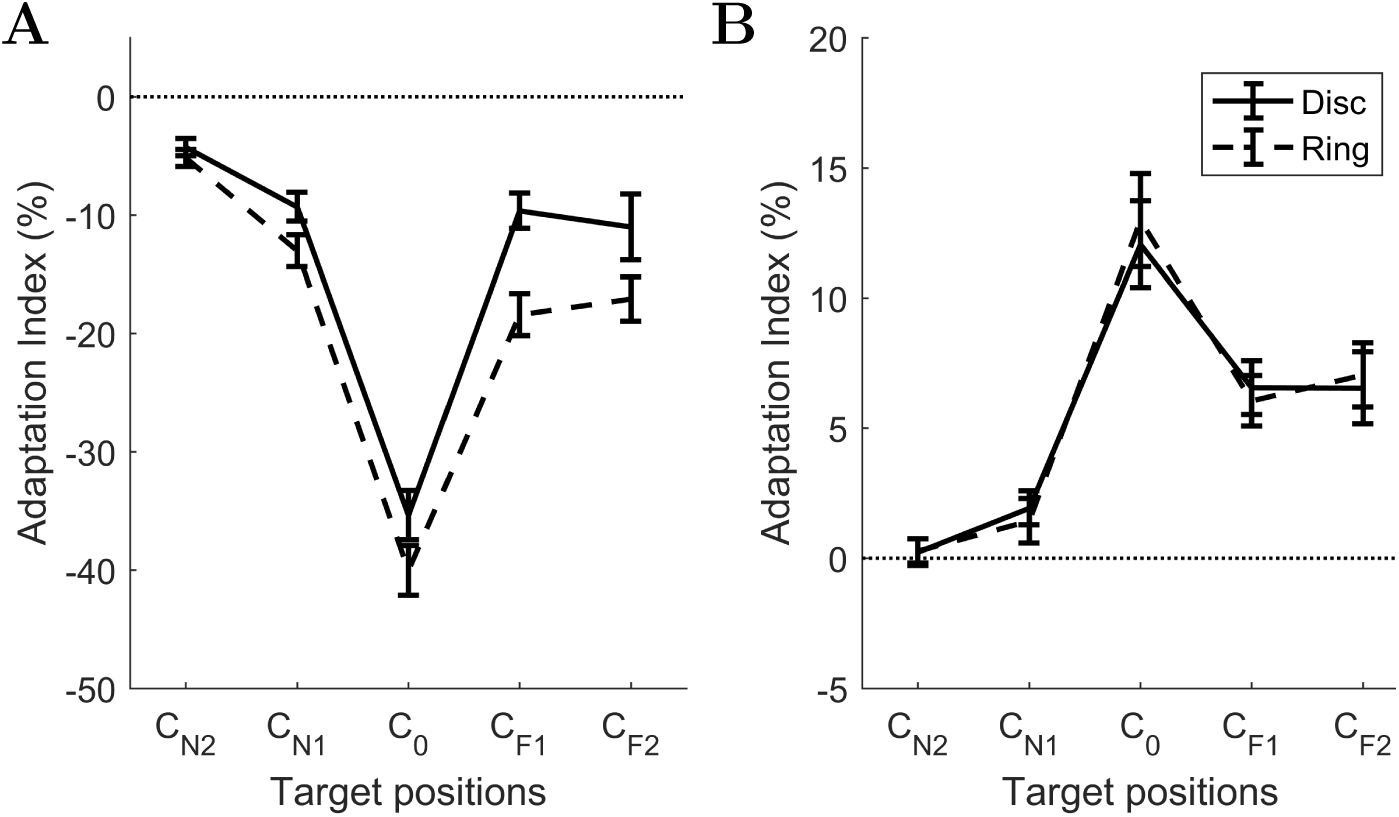
Results of Experiment 2 (dashed line) together with results from Experiment 1 (solid line). Average adaptation indices are shown as a function of five target positions, for small (A) and large (B) targets from both experiments. The pattern between the sets of results is remarkably consistent, showing that the adaptation effect does not critically depend on the geometry of the stimulus. Error bars: standard error of the mean (SEM). C: center, N2: nearer 2, N1: nearer 1, 0: zero, F1: further 1, F2: further 2.

Sizes of the small target rings presented at C_0_ were underestimated by 40% (SEM=2.1%) in the adaptation condition, as compared to the control condition. Such underestimations of size were also observed for other target positions. One sample *t* test analyses revealed that the AIs for small targets were significantly smaller than zero, at all of the five positions (FDR corrected *p*s < 0.001). Two-way repeated measures ANOVA showed that there was a significant main effect of target positions (*F*(4, 44) = 98.88, *p* < 0.001), but no significant main effect of adapter hemifield (*F*(1, 11) = 1.1371, *p* = 0.31).

Large target rings presented at C_0_ position were perceived 13% (SEM=1.8%) larger in the adaptation condition. Perceived sizes of large targets presented at the other 4 positions showed a similar overestimation of size, although with smaller magnitudes of effect. FDR corrected one sample *t* tests showed that the AI values for large targets were significantly higher than zero, at three of the five positions: 0, F1, and F2 (*p*s < 0.001). Moreover, repeated measures ANOVA revealed that there was a significant main effect of target positions (Sphericity was corrected via Greenhouse-Geiser correction; *F*(1.99, 21.99) = 29.99, *p* < 0.001), and no significant main effect of adapter hemifield (*F*(1, 11) = 4.39, *p* = 0.06).

Further analyses were conducted to compare the AIs along the central row obtained using the two types of stimuli. ANOVA results showed that there was a significant main effect of stimulus type for the small target size (*F*(1, 22) = 10.25, *p* = 0.004). In detail, AI was significantly smaller (*i*.*e*. the adaptation effect was greater) with small rings than with small discs. There was no significant interaction between the stimulus type and the target positions (sphericity corrected, *F*(2.38, 52.3) = 1.85, *p* = 0.16). For the large target sizes, there was no main effect of stimulus type on AI (*F*(1, 22) = 0.008, *p* = 0.93), and no interaction between stimulus type and target positions (sphericity corrected, *F*(2.34, 51.55) = 0.22, *p* = 0.84).

## 4 Discussion

The main purpose of this study was to systematically investigate the non-local effects of size adaptation in the visual field. In two behavioral experiments we measured the perceived sizes of small and large target stimuli (discs in Experiment 1, and rings in Experiment 2) in both left and right visual hemifields, following a period of adaptation on the same visual hemifield. Consistent with previous literature, our results showed that the adaptation to a mid-sized stimulus resulted in an underestimation of the size of subsequently presented small targets, and an overestimation of the size of large targets. Most importantly, the present study demonstrated that the effect of size adaptation is not spatially limited to the adapter’s location, instead, it spreads over a wide range of area in the visual field. Our results also showed that the adaptation effect does not critically depend on the exact geometry of the stimulus. Our findings agree with the previous studies on size adaptation (e.g. Pooresmaeili et al., 2013), and adds to the literature on the spatial extent of size aftereffect. Our data clearly showed that the effect was robust at both directly adapted and non-adapted positions with ring-shaped stimuli as well as filled disc stimuli.

Context-dependent changes in perceived size have been shown to be associated with the position shifts in the receptive fields (RFs) of neurons in the primary visual cortex, V1 (Ni, Murray, & Horwitz, 2014; He, Mo, Wang, & Fang, 2015). According to this model, an object may appear larger (smaller) owing to the context within which it is viewed, because the positions of RF centers of neurons that normally process the visual space corresponding to the stimulus and its near surround shift inward (outward), thus the stimulus is processed by a denser (sparser) array of neurons, which in turn leads to a larger (smaller) representation of the stimulus in the visual cortex. Could a similar mechanism account for the adaptation effect we found at the concentric position? It is unlikely because such a low-level model could not explain the effect at distant target positions, because RF shift model would predict either no or an opposite effect at these positions. The adaptation effect we observed at non-stimulated, distant positions implies the involvement of different mechanisms possibly including those in higher level visual areas.

The findings of Experiment 1 and 2 taken together further limits the possible underlying neuronal mechanisms. We found nearly identical patterns of results in Experiment 1, using solid discs, and Experiment 2, using rings. A simple model where activity of neuronal populations whose RFs fall within the boundaries of the solid discs are attenuated would not be sufficient to explain the results of both experiments. A model in which the size computation depends on the activity of edge units could potentially explain the results of both experiments simultaneously, but only at the adapter locations. Such a model would still fail to explain the results at distant locations.

Classic repulsive aftereffects are mostly thought to be a consequence of decreased sensitivity of “channels” that are selectively tuned to a particular stimulus feature, such as frequency or orientation (see Braddick, Campbell, & Atkinson, 1978; Blakemore, Nachmias, & Sutton, 1970). Based on this idea, a two-layer model can in principle explain the non-local size adaptation effect we found. In the model, units (or channels) in each layer are tuned to both size and location as shown in Figure 5. Output of Layer 1 units are fed forward to Layer 2 units. Critically, however, Layer 2 units pool signals from a number of Layer 1 units. Thus, effectively, Layer 2 units have wider tuning curves. For example, a certain size object at a certain position in space would activate only a few Layer 1 units, but because of pooling, a larger number of units in Layer 2, even those whose center of spatial tuning curves lie further away, may become active. Finally the output of Layer 2 units is sent to a higher level center that determines the perceived size and position based on the input and knowledge of tuning curves. When adapted to a certain size at a certain location, the sensitivity of the units that are responsive to that size and location decreases in both layers (Figure 5.B). But because the Layer 2 units have wider spatial tuning curves, their adaptation can influence the perceived size of objects presented even in parts of the visual field that was not stimulated by the adapter in the first place. A simple Gaussian spatial tuning curve in Layer 2 would also explain why the adaptation effect decreases as the distance between the adapter and target stimuli increases. Here, in this conceptual model we only considered two layers and a feedforward relationship between the two layers. Of course it is also possible that there are more than two layers with a more complex interaction, including feedback messaging among the layers. Note that, other alternatives to the simple gain control mechanism suggested in the model (i.e. channel responses decreasing with adaptation) can easily be incorporated, such as the recently proposed models of adaptation based on a normalization model where weights of interaction between units are updated based on an Hebbian rule during adaptation (Carandini & Heeger, 2012; Yiltiz, Heeger, & Landy, 2020). In those models, adaptation depends on not only the recent history of a single neuron (or channel) response, but the history of a pool of neurons, which is consistent with our findigns.

**Figure 5:**
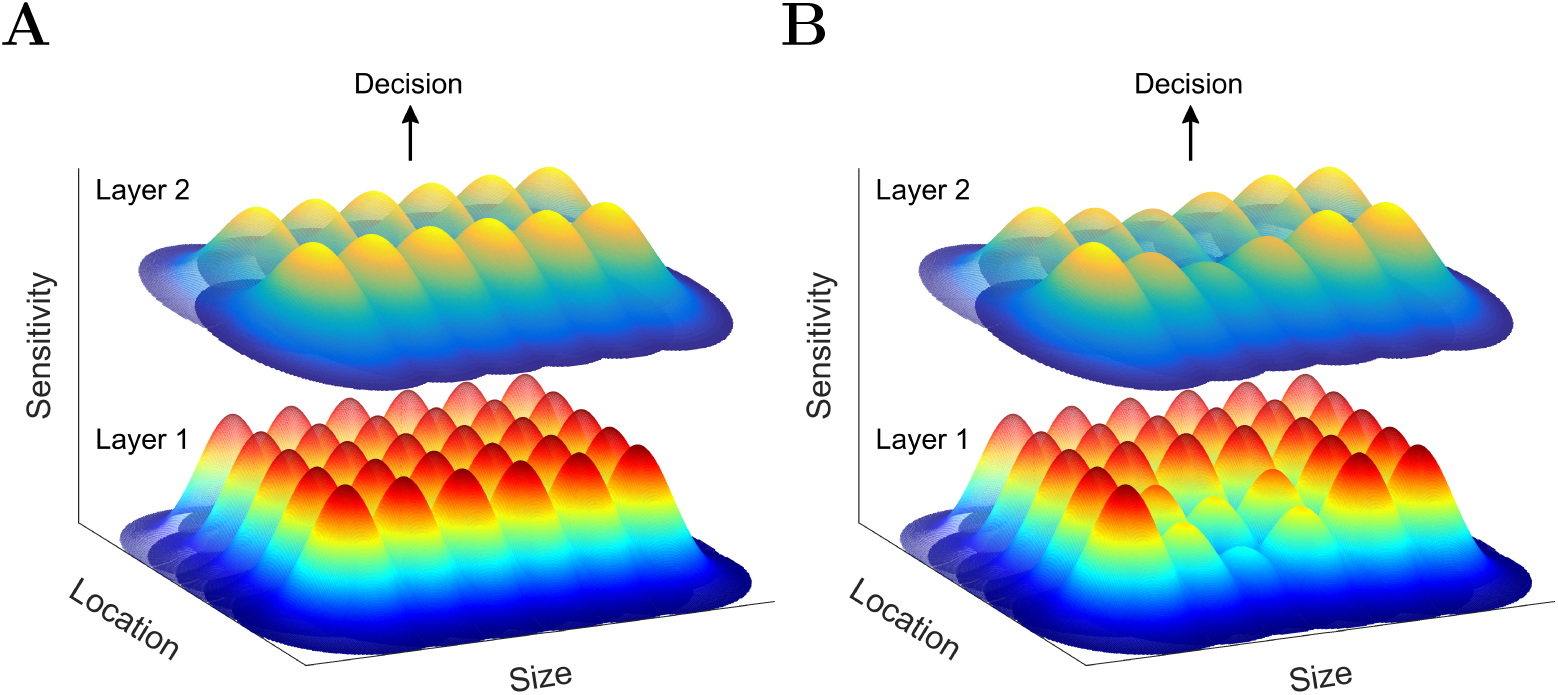
A possible explanation for the size aftereffect at the eccentric positions. A. The size and location tuned channels without any adaptation. B. Altered sensitivity profiles of channels after adaptation to a size at a certain location. Two layers of size channels are shown here. The lower layer consists of narrower channels which are responsive to a smallest range of sizes at very limited area. The higher layer receives input from the first layer, and responsive to a wider range of sizes with a greater width.

Is such a simple computational model biologically plausible? Pooresmaeili et al., 2013 studied the effect of adaptation using fMRI and found that the activated area of cortical surface in V1, as well as V2, V3 and V4 (although not statistically significantly in V4) correlated with the perceived size (but see Chouinard & Ivanowich, 2014, for a critical review). There are also studies supporting the feedback modulation on size adaptation effect. For example Kreutzer, Fink, and Weidner (2015) reported a modulatory effect of attention on the effect of size adaptation, signalling the involvement of higher level visual areas. In line with this finding, Laycock et al. (2017) reported that the adapter does not affect subsequent size perception when the participants are not consciously aware of the adapter stimulus. These results support a neural mechanism of size perception that relies on responses in multiple visual areas, as proposed by the computational model introduced above.

In both of the experiments, we observed an asymmetry between the adaptation indices of small and large targets. The magnitude of effect for small targets almost triples that for large targets. Such an asymmetry between size under- and overestimation was also present in previous studies (Pooresmaeili et al., 2013; Schwarzkopf & Rees, 2013). This might be due to the difference between the amounts of information available from the two sizes of target stimuli. Suzuki and Cavanagh (1998) showed that the magnitude of shape aftereffect was inversely linked to the magnitude of the signal available from the target stimulus. When the image signal in the test phase was abundant (e.g. when the duration of the test phase was long, or the luminance contrast of the target stimulus was high) the adaptation effect was weaker. They suggested that this is because higher image signal helps visual system to correct the initial biased perception. Based on this correction hypothesis, large target stimuli in our study might be providing further signal to the system as compared to the small targets, so that the magnitude of the adaptation effect for the large targets is weaker (i.e. corrected to a greater degree) than that for the small targets. Relatively weaker adaptation effect that was observed near fixation point (See Figure 3) can also be explained with such a correction hypothesis. It is known that the density of ganglion cells and visual sensitivity is maximum near fovea and decreases as the eccentricity increases (Wässle, Grünert, Röhrenbeck, & Boycott, 1990; Rijsdijk, Kroon, & van der Wildt, 1980). Therefore it could be that we observe less illusory effect near the fixation point given that the amount of information that the visual system receives would be abundant for the areas near fixation point, as the correction hypothesis would predict. Alternatively, it is also possible that the participants shifted their strategy for nearer positions. Although the participants were instructed to make their decision according to the stimuli size only, distance (e.g. distance from the fixation point) information was available as an additional reliable cue when the stimuli were presented in the nearer locations.

Another noteworthy finding was the asymmetry between two visual hemifields. Previous studies usually investigated size adaptation effect only in one visual hemifield (e.g. Pooresmaeili et al., 2013; Zeng et al., 2017; Zimmermann, Morrone, & Burr, 2016). Here we showed that the magnitude of the adaptation effect may differ depending on the visual hemifield in which the adapter was presented. Although not consistently significant, we found greater adaptation effects on the left visual hemifield, which suggests a hemispheric asymmetry. There are, however, conflicting findings in literature about visual field asymmetries when it comes to size judgments. For instance, Muller-Lyer illusion has been found to be stronger in the left visual hemifield (Clem & Pollack, 1975). Conversely, Saneyoshi (2018) found greater size overestimation effect with Ebbinghaus inducers on the right visual field. Also size discrimination has been found to be more accurate in the left visual field, implying that the perceived size is more resistant to illusory effects in the left visual field (Corballis, Funnell, & Gazzaniga, 2002). Nevertheless, it is more likely that the asymmetry is a consequence of a particular combination of many different stimulus properties such as luminance, eccentricity and exposure duration as it has been shown that all features of visual input contribute to the hemispherical superiority via complex interactions (Sergent, 1983).

Previous studies have shown that the adaptation phenomenon have several benefits for the visual system. It increases the salience of a novel stimulus (McDermott, Malkoc, Mulligan, & Webster, 2010), leads to an enhancement in the ability to discriminate a particular feature of a stimulus (Greenlee & Heitger, 1988), and maximizes the efficiency by serving as a gain control mechanism (Wainwright, 1999; Barlow, 1990). Naturally, a mechanism where these favorable effects are not tied to the exact retinotopic position would make a more efficient system. Our results support that indeed this happens in the visual system. Our findings may have further implications on how the visual system maximizes the efficiency and how the visual information is being encoded. These issues and the underlying mechanism of the non-local size adaptation effect should be modelled rigorously (not only conceptually) in a future work, where neuronal responses are also investigated.

## 5 Conclusion

To conclude, we demonstrated that prolonged exposure to a certain size stimulus distorts the subsequent size perception within a wide spatial extent, not limited to the position of the adapter. Overall results of two experiments suggest that neither a model in which neural receptive fields shift, nor a mechanism in which neuronal activity is simply attenuated with prolonged exposure to an adapter, is adequate to explain the non-local adaptation effect. A simple multi-layered computational model, however, can parsimoniously explain the non-local distortion in perceived size following adaptation.

## 6 Author Contributions

EA and HB conceived the original idea. EA designed, implemented and conducted the experiments, and analysed the data. EA and HB wrote the manuscript.

## 7 Declaration of Interest

Declarations of interest: none

## 8 Acknowledgments

